# A 16S rRNA gene-based analysis of microbial communities in compost-bedded pack barns from dairy farms in Argentina

**DOI:** 10.64898/2026.04.04.716490

**Authors:** Juan L. Monge, Cecilia Peralta, Leopoldo Palma

## Abstract

Microbial communities play a central role in compost-bedded pack (CBP) systems by driving organic matter decomposition and nutrient cycling. The objective of this study was to characterize and compare the bacterial community structure of CBP from two dairy farms in Córdoba, Argentina, using 16S rRNA gene sequencing. Two CBP systems were evaluated: Martin Bono (MB; 30 months in operation) and Angela Teresa (AT; 20 months). The MB system was established on natural soil without bedding addition and included concrete feed alleys, whereas AT was initiated with peanut shell bedding and lacked concrete alleys. In both systems, compost was tilled twice daily. Two samples per farm were collected at a depth of 30 cm during winter 2019. Raw Illumina reads were processed using the DADA2 pipeline, including quality filtering, error modeling, denoising, and chimera removal. A total of four samples yielded 2,503 amplicon sequence variants (ASVs), with approximately 76% of reads retained after filtering and chimera removal, indicating high-quality sequencing data. Taxonomic analysis revealed that bacterial communities in both systems were dominated by phyla typically associated with compost environments, including Actinobacteriota, Proteobacteria, and Firmicutes. Differences in relative abundance between systems suggested shifts in community composition associated with management conditions.

## Introduction

Compost-bedded pack (CBP) systems are increasingly adopted in dairy production due to their benefits in animal welfare, cow comfort, and manure management efficiency (Biasato et al., 2019; Black et al., 2013). In these systems, microbial communities play a central role in organic matter decomposition, heat generation, and nutrient cycling, driving the biological processes that sustain compost functionality (Insam and de Bertoldi, 2007; Sánchez-Monedero et al., 2001). Nitrogen transformations mediated by microorganisms are critical for maintaining compost quality and environmental sustainability, as they regulate ammonia volatilization, nitrate formation, and overall nitrogen balance (Kumar et al., 2025).

The structure and function of microbial communities in CBP systems are strongly influenced by management factors such as bedding material, aeration, moisture content, stocking density, and system age (Black et al., 2013; Endres and Barberg, 2007). These variables shape physicochemical conditions within the pack, thereby affecting microbial activity, diversity, and metabolic pathways. Among microbial functional groups, nitrifying bacteria are of particular interest, as they mediate the oxidation of ammonia to nitrite and nitrate, playing a key role in nitrogen cycling and influencing nitrogen losses through volatilization and leaching (Kowalchuk and Stephen, 2001; Prosser et al., 2007).

Previous studies have reported the presence of highly diverse microbial communities in composting environments, including taxa associated with organic matter degradation and nutrient transformations (Kumar et al., 2025; Palaniveloo et al., 2020). However, limited information is available on how CBP management practices specifically influence the abundance and diversity of nitrifying bacteria in dairy production systems. This represents an important knowledge gap, given the relevance of nitrogen cycling for both compost efficiency and environmental impact.

Advances in high-throughput sequencing technologies, particularly 16S rRNA gene amplicon sequencing, have enabled detailed characterization of microbial communities and their potential functional roles in complex environments such as compost-bedded packs (Callahan et al., 2016; Knight et al., 2018). These approaches allow for the identification of key microbial taxa and provide insights into how management practices shape microbiome composition.

The objective of this study was to characterize and compare the bacterial community structure of CBP systems from two dairy farms in Córdoba, Argentina, with particular emphasis on nitrifying bacteria, using 16S rRNA gene sequencing.

## Materials and Methods

### Study sites and sampling

Two compost-bedded pack (CBP) dairy systems located in Córdoba, Argentina, were evaluated: Martin Bono (MB; 30 months in operation) and Angela Teresa (AT; 20 months). The MB system was established on natural soil without bedding addition and included concrete feed alleys, whereas AT was initiated with peanut shell bedding and lacked concrete alleys. In both systems, compost was tilled twice daily.

Sampling was conducted during winter (July 2019). Two samples were collected from each CBP system at a depth of approximately 30 cm, resulting in a total of four samples (AT1, AT2, MB1, and MB2).

### DNA extraction and sequencing

Soil samples were first lyophilized to facilitate handling and homogenization due to their sticky consistency. Subsequently, dried samples were homogenized in a high-speed mixer (High-speed Universal Disintegrator, Pro-Lab Diagnosis) and DNA extraction was performed using the microbiome DNA purification kit PureLink™ (ThermoFisher) and subjected to 16S rRNA gene amplicon sequencing using Illumina technology (INDEAR, Argentina), following standard protocols for microbiome characterization (Caporaso et al., 2012). Paired-end reads were obtained and merged prior to downstream analysis.

### Bioinformatic processing

Raw sequence data were initially processed in Geneious Prime (version 2025.1.3), where trimming and quality filtering were performed to remove low-quality reads and adapter sequences. Subsequently, the filtered reads were analyzed using the DADA2 pipeline in R (Callahan et al., 2016). This included error rate learning, sequence denoising, and inference of amplicon sequence variants (ASVs). Chimeric sequences were identified and removed using a consensus-based approach.

After processing, a total of 2,503 amplicon sequence variants (ASVs) were obtained across four samples, with approximately 76% of reads retained after filtering and chimera removal. Taxonomic classification of ASVs was performed using the SILVA reference database (v138.1) (Quast et al., 2013), assigning taxonomy across multiple levels, including phylum and genus.

### Statistical and ecological analysis

Microbial community analyses were conducted using the (McMurdie and Holmes, 2013). Relative abundance of taxa was calculated and visualized at the phylum level. Alpha diversity was assessed using the Shannon index (Shannon, 1948). Beta diversity was evaluated using Bray– Curtis dissimilarity (Bray and Curtis, 1957), and principal coordinate analysis (PCoA) was used to visualize differences in community structure between systems.

Nitrifying bacteria were identified based on taxonomic assignment at the genus level, focusing on *Nitrosomonas, Nitrosococcus*, and *Nitrobacter*. Differences in abundance and diversity between systems were evaluated descriptively due to the limited number of replicates. Additionally, genera associated with mastitis-related pathogens, including *Staphylococcus, Streptococcus, Corynebacterium*, and *Escherichia*, were specifically examined to assess potential sanitary risks.

## Results

### Microbial diversity and community structure

Alpha diversity, estimated using the Shannon index, showed slightly higher values in MB compared to AT samples (Figure 1). However, these differences were not statistically significant (Wilcoxon test, *p* = 0.33), likely due to the limited number of replicates. Despite this, MB samples consistently exhibited higher diversity values, suggesting a trend toward increased microbial complexity.

**Figure 1.**
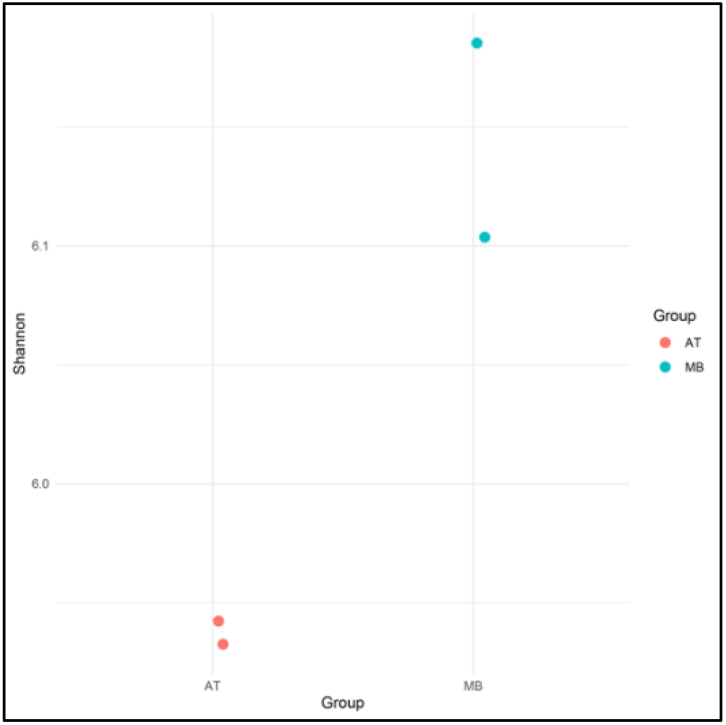
Shannon diversity index of bacterial communities in compost-bedded pack systems (AT and MB). Boxplots represent alpha diversity values for each system. No significant differences were observed between groups (Wilcoxon test, p = 0.33).

Beta diversity analysis based on Bray–Curtis dissimilarity revealed a clear separation between AT and MB samples (Figure 2), indicating distinct microbial community structures between systems. Replicates clustered according to treatment, supporting the reproducibility of the observed patterns.

**Figure 2.**
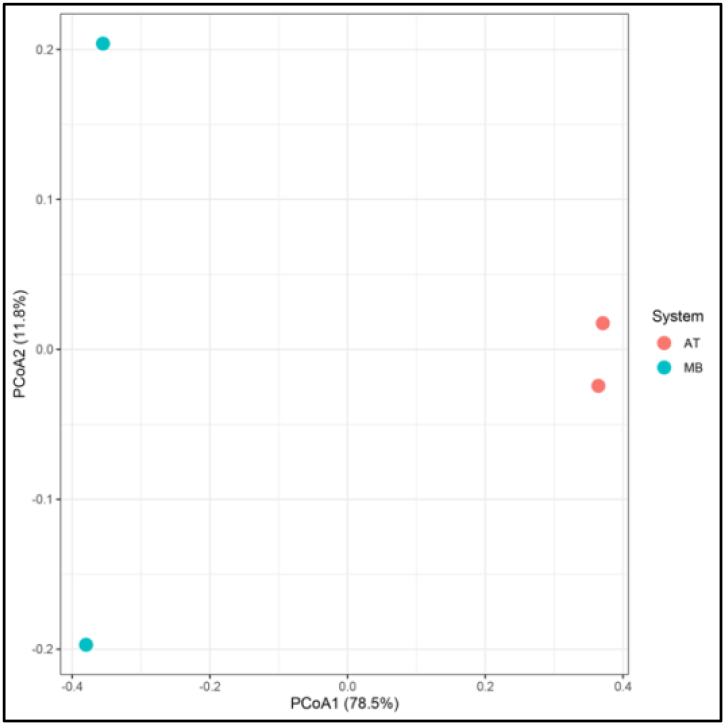
Principal coordinate analysis (PCoA) based on Bray–Curtis dissimilarity showing differences in bacterial community structure between compost-bedded pack systems (AT and MB). Each point represents one sample, and colors indicate system type. The first principal coordinate explains 78.5% of the variation.

### Mastitis-associated genera

Genera associated with mastitis-related pathogens were detected in both systems, including *Corynebacterium, Pseudomonas*, and *Staphylococcus* (Figure 3).

**Figure 3.**
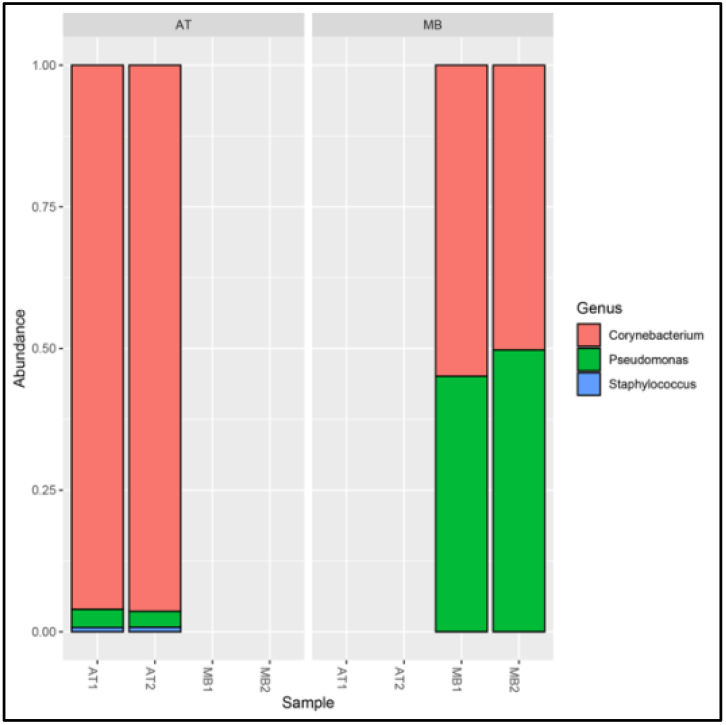
Relative abundance of mastitis-associated genera (*Corynebacterium, Pseudomonas, Staphylococcu*s) in AT and MB samples.

AT samples were dominated by *Corynebacterium*, with only minor contributions from other genera. In contrast, MB samples showed a more balanced composition, with increased relative abundance of *Pseudomonas* and detectable levels of *Staphylococcus*. These results suggest differences in the potential sanitary profile between systems.

### Nitrifying bacteria

Nitrifying bacteria were detected at the genus level, including *Nitrosomonas* and *Nitrosococcus* (Figure 4). These taxa were present in both systems, although their relative abundance appeared higher and more consistent in MB samples, suggesting enhanced nitrogen cycling potential in this system.

**Figure 4.**
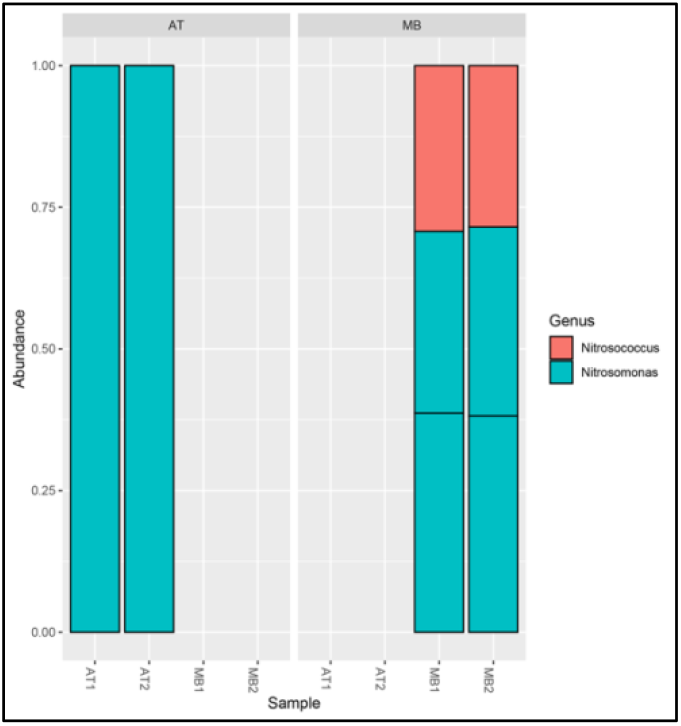
Relative abundance of nitrifying bacteria (*Nitrosomona*s and *Nitrosococcus*) across samples from both CBP systems.

### Functional microbial groups

Analysis of key functional genera associated with compost processes revealed marked differences between systems (Figure 5). AT samples were largely dominated by *Pseudomonas*, indicating a simpler functional structure. In contrast, MB samples displayed a broader functional diversity, including the presence of *Mycobacterium* and *Streptomyces*, in addition to nitrifying bacteria such as *Nitrosomonas*.

**Figure 5.**
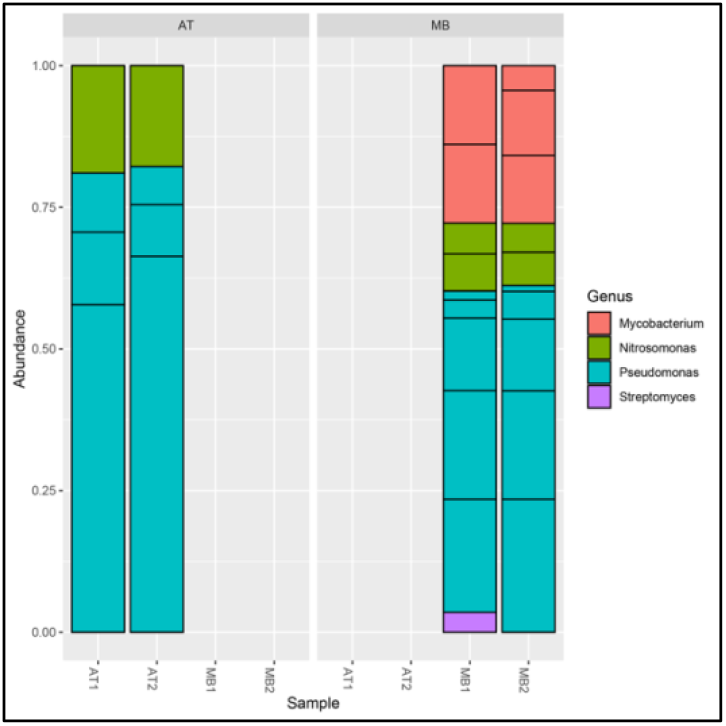
Relative abundance of selected functional genera associated with compost processes (*Mycobacterium, Nitrosomonas, Pseudomonas, Streptomyces*) in AT and MB systems.

The exclusive detection of *Mycobacterium* in MB samples further highlights differences in environmental conditions and microbial niches between systems. Overall, these findings suggest a more complex and functionally diverse microbial community in MB.

### Community abundance patterns

Heatmap analysis of the most abundant genera revealed distinct community patterns between systems (Figure 6). AT samples exhibited a more homogeneous microbial profile, with higher relative abundance of *Corynebacterium*, whereas MB samples showed a more heterogeneous distribution of taxa, including increased representation of several genera associated with compost processes. These patterns further support differences in microbial structure between CBP systems.

**Figure 6.**
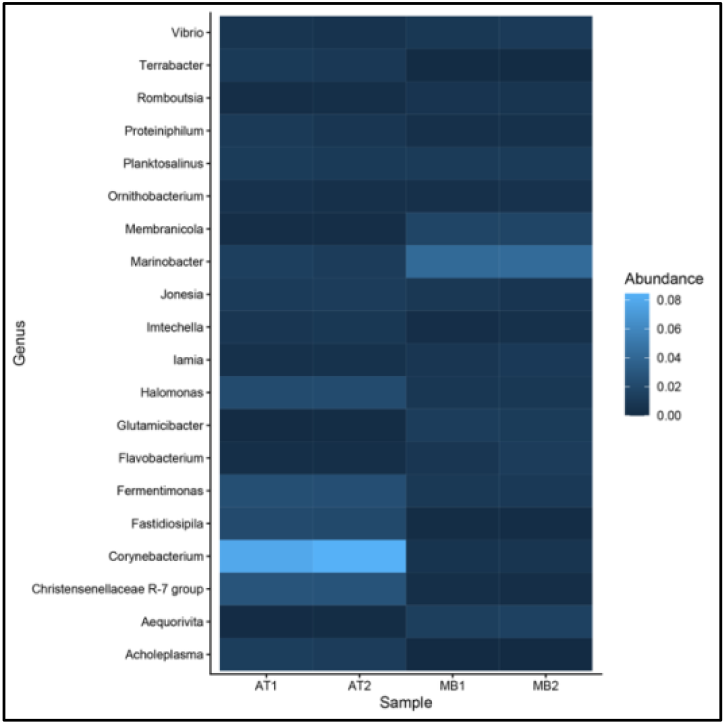
Heatmap showing the relative abundance of the most abundant bacterial genera across samples. Color intensity represents relative abundance, highlighting differences in microbial composition and heterogeneity between AT and MB systems.

## Discussion

The present study provides a first 16S rRNA gene-based characterization of microbial communities in compost-bedded pack (CBP) barns from dairy farms in Córdoba, Argentina. Although based on a limited number of samples, the results show that bacterial community composition differed clearly between the two evaluated systems, suggesting that management conditions, bedding characteristics, and system configuration can strongly influence microbial structure in CBP environments.

Both systems were dominated by phyla commonly associated with composting and organic matter degradation, including Actinobacteriota, Proteobacteria, and Firmicutes. This is consistent with the biological function of CBP systems, where microbial activity supports decomposition, heat generation, and nutrient turnover (Insam and de Bertoldi, 2007; Tiquia et al., 2002). The separation observed in the Bray–Curtis PCoA indicates that, despite the shared productive context, each barn harbored a distinct bacterial assemblage. The close clustering of replicates within each farm further supports the consistency of the microbial profiles detected.

The MB system showed a tendency toward higher alpha diversity and a broader distribution of taxa compared with AT. This pattern may reflect the combined influence of longer operational time, absence of initial bedding addition, and the presence of concrete feed alleys, all of which may affect moisture gradients, organic matter inputs, and aeration dynamics (Endres and Barberg, 2007; Endres and Janni, 2008). In compost-based systems, greater microbial diversity is often associated with increased functional redundancy and process stability, which may be advantageous for maintaining compost performance under variable environmental conditions (Allison and Martiny, 2008; Shade et al., 2012). However, because of the small sample size, these differences should be interpreted cautiously.

Particular attention was given to nitrifying bacteria because of their relevance to nitrogen transformations in compost systems. Genera such as *Nitrosomonas* and *Nitrosococcus* were detected in both farms, but their abundance appeared greater and more consistent in MB. This suggests that microbial processes linked to ammonia oxidation may be favored under the environmental and management conditions present in that system. Since nitrogen cycling is a central component of compost quality and environmental performance, these differences may have practical implications for ammonia retention, nitrate formation, and overall nutrient dynamics in CBP barns (Kowalchuk and Stephen, 2001; Sánchez-Monedero et al., 2001). Nonetheless, functional activity was inferred only from taxonomic composition, and future studies should incorporate direct measurements of nitrogen transformation rates or functional genes involved in nitrification.

The detection of genera associated with mastitis-related pathogens, including *Staphylococcus, Streptococcus, Corynebacterium*, and *Escherichia*, also highlights the sanitary relevance of microbial monitoring in these systems. Although 16S rRNA sequencing does not allow confirmation of pathogenic strains or viability, the presence and relative abundance of these genera suggest that CBP management may influence microbial groups with potential relevance to udder health (Bradley, 2002; Smith et al., 1985). Differences between systems may therefore reflect not only composting performance but also distinct sanitary profiles linked to bedding composition, moisture, and aeration

This study has some limitations that should be acknowledged. The analysis was based on only four samples collected at a single time point during winter, which limits statistical power and prevents evaluation of seasonal dynamics. In addition, taxonomic inference from 16S rRNA amplicons provides limited functional resolution (Janda and Abbott, 2007; Knight et al., 2018). Even so, the results offer a useful initial description of microbial community patterns in Argentine CBP systems and identify management-associated differences that merit deeper investigation. Future work should include a larger number of farms, repeated sampling over time, and integration of physicochemical variables such as temperature, moisture, pH, and nitrogen forms to better explain microbial shifts and their functional consequences.

In conclusion, the bacterial communities of the two CBP systems differed in both composition and inferred functional potential. The MB system showed higher diversity and a greater representation of nitrifying bacteria, suggesting that barn design and management history may shape key microbial processes in compost-bedded pack environments. These findings contribute to a better understanding of microbiological dynamics in dairy CBP systems and may support future improvements in compost management, nutrient cycling, and animal health.

## Author contributions

Conceptualization, L.P., methodology, J.M., C.P. and L.P.; formal analysis, J.M., C.P. and L.P.; investigation, J.M.; C.P. and L.P.; resources, J.M. and L.P.; data curation, L.P.; writing—original draft preparation, L.P.; writing, review and editing, L.P.; visualization, L.P.; supervision, J.M. and L.P.; project administration, J.M. and L.P.; funding acquisition, J.M. and L.P. All authors have read and agreed to the published version of the manuscript.

## Acknowledgments

Leopoldo Palma gratefully acknowledges the Spanish Ministry of Science, Innovation, and Universities, the Spanish State Research Agency, and the European Union for funding his Ramón y Cajal contract (grant ref. RYC2023-043507-I).

## Data availability

The raw sequencing reads have been deposited in the NCBI Sequence Read Archive (SRA) under BioProject accession PRJNA1449009, with associated BioSample accession numbers XXX.

